# Gamma–Theta–Spike Coupling Coordinates Sequential Memory in Human MTL

**DOI:** 10.1101/2025.06.24.661371

**Authors:** Muthu Jeyanthi Prakash, Johannes Niediek, Rainer Surges, Florian Mormann, Stefanie Liebe

## Abstract

It has been suggested that hierarchical synchronization of theta and gamma oscillations coordinates neural activity during sequence memory. Yet, the role of gamma oscillations and their interaction with theta and single-unit activity (SUA) has not been directly examined in humans. We analysed simultaneous micro wire recordings of single-unit activity (N = 1417) and local field potentials (N = 917 channels) from the medial temporal lobe (MTL) of epilepsy patients performing a visual multi-item sequence memory task. During encoding, both spiking activity and gamma power contained item-specific information and were temporally coupled. During memory maintenance, stimulus-specific gamma was characterized by recurring bursts during which spiking was tightly synchronized and both, gamma and spiking, were preferentially aligned to similar theta phases predictive of sequential stimulus position. These findings demonstrate that theta–gamma–spike interactions support a phase-based multiplexed code for sequential memories in the human MTL.

## 1 Introduction

Theta (2–8 Hz) and gamma (30–100 Hz) oscillations are fundamental to memory processing across species [21, 4]. Their hierarchical synchronization has been proposed to temporally structure spiking activity via a nested, multiplexed theta–gamma–spike code [15, 11, 16, 5, 21]. The medial temporal lobe (MTL) is a key structure for learning and memorizing sequential information [13, 19, 2]. A recent study provided evidence for a theta-dependent phase-of-firing code for sequence position during memory maintenance within the MTL [14]. However, how gamma oscillations interact with theta rhythms and single-unit activity during sequence memory remains unresolved. To address this question, we analysed micro-wire recordings of local field potentials (LFPs) and SUA from the MTL of epilepsy patients performing a visual sequence memory task. Our study reveals several novel insights into how gamma oscillations participate in organizing spiking activity within theta cycles to support memory for stimulus sequences. During encoding, both gamma power and single-unit activity carried stimulus-specific information, but selectivity in gamma emerged earlier than in spiking activity, despite being overall less discriminative. During memory maintenance, gamma activity reappeared in burst-like patterns, and stimulus-specific increases in firing rates were confined to these bursts. Moreover, both single-unit activity and gamma power exhibited position-dependent phase coupling to theta oscillations. Notably, gamma and spiking activity were aligned to similar theta phases, indicating a shared temporal reference across signals. Together, these findings suggest that coordinated theta–gamma–spike interactions structure the temporal organization of sequential memory content in the human MTL.

## 2 Results

Task and recording details are described in detail in [14]. In brief, we recorded local field potentials (917 channels) and spiking activity (1417 units) from the hippocampus (HPC), entorhinal cortex (EC), amygdala (AM), and parahippocampal cortex (PHC) of epilepsy patients undergoing pre-surgical monitoring while they performed a visual sequence memory task (Fig.1a). In each trial, participants viewed a sequence of four images (200 ms each, 200 ms inter stimulus interval), followed by 2500 ms delay and then chose the correct order from a probe panel (Fig.1b).

**Fig. 1.**
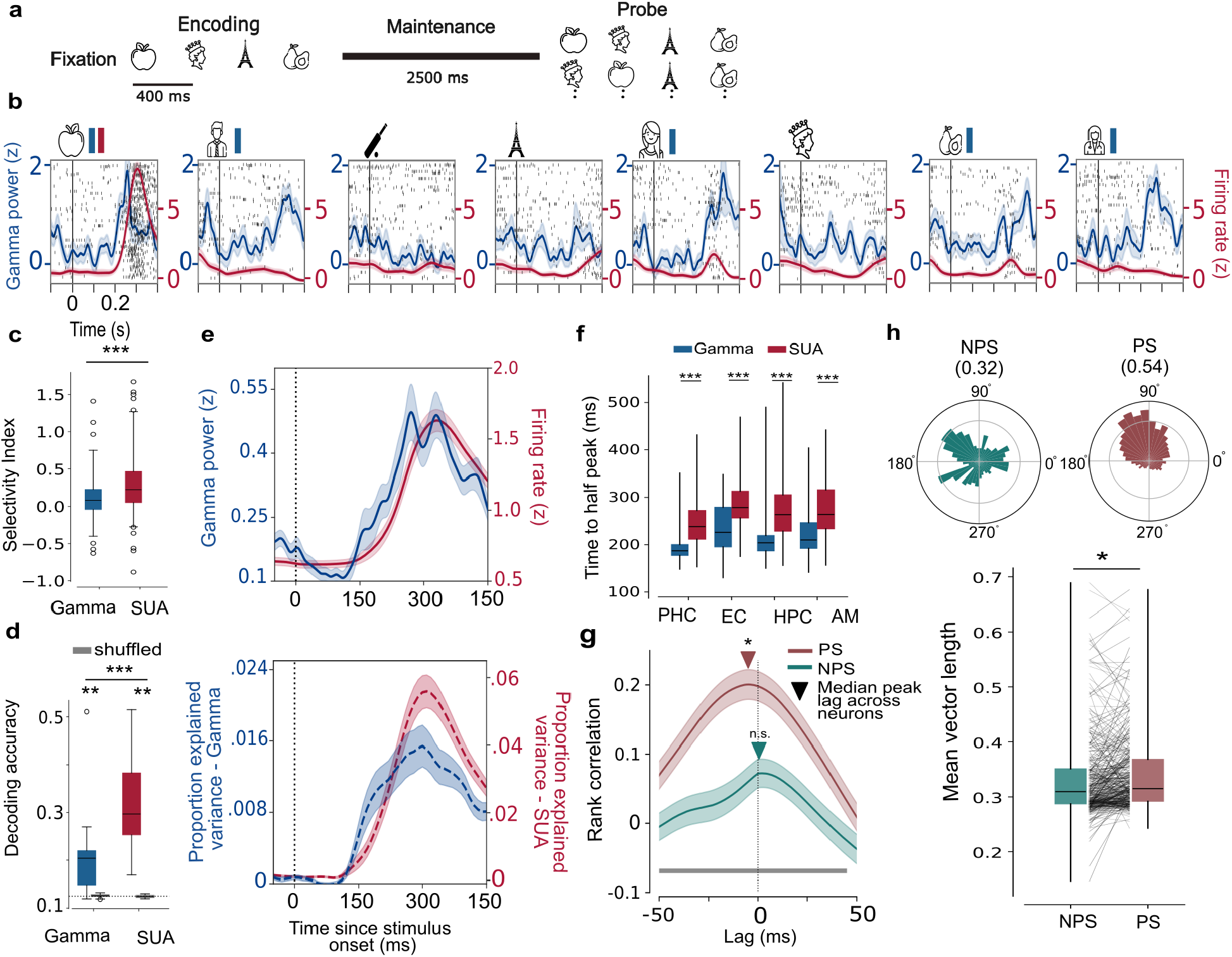
Relationship between spiking and gamma during encoding. **a**, Structure of the multi-item sequence memory task performed by participants. **b**, Firing rate and Gamma power (*γ*, 60 - 100 Hz) z-scored with respect to baseline are plotted along with raster for all stimuli shown in a session. Vertical line denotes time of stimulus onset. The gamma power and neuron recorded from the same channel, showed the strongest response to the first stimulus (preferred stimulus, PS, p *<*0.001, one - sided Wilcoxon signed-rank (WSR) test). Red and blue bar near stimulus denotes significant increase in firing rate and gamma power, respectively following stimulus onset. **c**, Selectivity index (Hedge’s g between response to first and second strongest response eliciting stimulus) for Gamma power and SUA, p *<*0.001, one-sided Mann Whitney U test. **d**, Decoding accuracy for stimulus-identity (proportion correct) using average gamma power (60-100 Hz) and firing rates in responsive channels / neurons as features. Dashed lines corresponds to performance expected by chance. (1/8) and performance using shuffled identity labels are shown in gray, p *<*0.001 one-sided Mann Whitney U test **e**,Temporal evolution of z-scored to baseline gamma power (*γ*, 60 - 100 Hz) and firing rate following presentation of the PS (top). Proportion Explained Variance in the gamma power and firing rates by the stimulus identity across time (bottom). Note different axis scaling for SUA vs gamma. **f**, Median latencies following stimulus onset for Gamma (Blue) and SUA (Red) for four MTL regions. p *<*0.001, Wilcoxon rank sum test (N = 416). **g**, Cross-correlogram between trial-averaged firing rates and Gamma power following PS and NPS onset. The gray bar at the bottom denotes significant difference between PS and NPS correlation (p *<*0.05, two-sided Permutation test, N = 416). Gamma precedes spiking during PS but not NPS trials (* p *<*0.05, n.s. non-significant, one-sided WSR test). **h**, Circular gamma phase-Histogram for a representative neuron shows temporal coupling following PS as opposed to NPS presentation during encoding. (Bottom) Spike phase coupling magnitude in stimulus encoding neurons is higher following PS vs. NPS presentation (N = 356, p *<*0.05, one-sided WSR test)

First, we established whether gamma oscillations in MTL encode stimulus information and asked whether the stimulus selectivity of gamma and SUA are comparable in human MTL as has been suggested for cortical regions previously [1]. In total, 31 percentage of channels (N = 291) exhibited significant stimulus-evoked increases in gamma power (60–100 Hz) relative to baseline (Wilcoxon Signed Rank (WSR) test, p *<* 0.05, Proportion of channels - HPC: 0.4; AM: 0.26; PHC: 0.23; EC: 0.08). 30 percentage of neurons (N = 422) encoded stimulus information (WSR test vs baseline, p *<* 0.05, proportion of units - HPC: 0.35; AM: 0.26; PHC: 0.21; EC:0.16). In accordance, both signals exhibited stimulus responses and were discriminative between different stimuli. Following the onset of a neuron’s preferred stimulus, gamma power increased significantly in channels with responsive neurons, both in magnitude (p *<* 0.001, Wilcoxon Rank Sum (WRS) test, z=7.96, Cohen’s d=0.57) and stimulus specificity, as reflected by greater explained variance (PEV) for stimulus identity (WRS test, z=8.4, p *<* 0.001, Cohen’s d=0.45; Fig.1e bottom). SUA demonstrated significantly higher selectivity, as measured using between the best response eliciting (preferred stimulus, PS) and second-best preferred stimulus compared to gamma power (Hedges g, p *<* 0.001, Mann-Whitney U test; Fig.1c). Similarly, stimulus decoding accuracy was significantly higher using firing rates, but could be decoded significantly above chance from gamma power. Interestingly, gamma systematically preceded firing rate increases (Fig.1e) for both response magnitude and stimulus information across all MTL regions (Median response time difference (Gamma - SUA); HPC: 67 ms; AM: 63 ms; PHC: 62 ms; EC: 71 ms; p *<* 0.001, WRS test, Fig.1f). A cross-correlation analysis also showed significantly higher correlation during PS than NPS presentation (p *<* 0.05, two-sided Permutation test, N = 416) and was characterized by a negative peak lag, pointing to the precedence of gamma relative to SUA (p *<* 0.05, WSR test, Fig.1g). Next, we assessed whether the observed stimulus-dependent co-modulation in gamma power and firing rate was accompanied by a stronger temporal coupling between both signals. Assessing spike–gamma coupling – using mean vector length with resampling to account for spike count differences – we indeed observed significantly stronger coupling following PS vs. NPS onset (one-sided WSR test, p *<* 0.05; Fig.1h (bottom)). In summary, our analyses show that LFP-gamma activity encodes stimulus information similar to SUA, albeit with an earlier onset and lower selectivity. This is accompanied by stronger temporal coupling between, suggesting a tight stimulus-dependent co-modulation in both magnitude and time during visual encoding in human MTL (p *<* 0.001, Mann-Whitney U test Fig.1d).

During memory maintenance increases in gamma magnitude have been proposed to represent the reactivation of previously encoded stimuli [17]. In line with that, on average LFP channels that encoded stimuli exhibited increased gamma power during the memory period (p *<* 0.001, Mann Whitney U test). However, we did not observe a significant difference in *average* gamma power across maintenance between PS and NPS (p *>* 0.05, WSR test). Instead, we did observe periods of significantly increased gamma power that reoccurred in a temporally structured manner throughout the delay as tested by cluster-based permutation tests (p *<* 0.05). To examine the functional relevance of observed gamma bursts, we identified significant bursts associated with each neuron’s preferred (PS) and non-preferred stimulus (NPS, p *<* 0.05, one-sided cluster-based permutation t-test). While overall burst frequency and duration did not differ between PS and NPS trials (p *>* 0.05, permutation t-tests), the increase in gamma power was significantly higher during PS-associated bursts, and neurons exhibited increased firing during PS-bursts (p *<* 0.001, one sided permutation test, N= 340; Fig. 2f). Randomly shifting burst windows abolished all effects, further supporting a stimulus-specific increase in firing. During bursts, we did not observe stronger spike–gamma phase locking, though this may reflect reduced phase estimation reliability because of lower firing rates during non-burst segments. Overall, our results demonstrate the occurrence of stimulus-specific gamma bursts during memory maintenance, which suggests that temporally structured oscillatory activity in the human MTL reflects encoded information in working memory. The fact that bursts coincide with elevated spiking implicates gamma oscillations in structuring neural activity during memory maintenance. This could be in line with the suggestion that gamma bursts reflect brief windows of increased neuronal excitability during multi-item memory [17, 15].

**Fig. 2.**
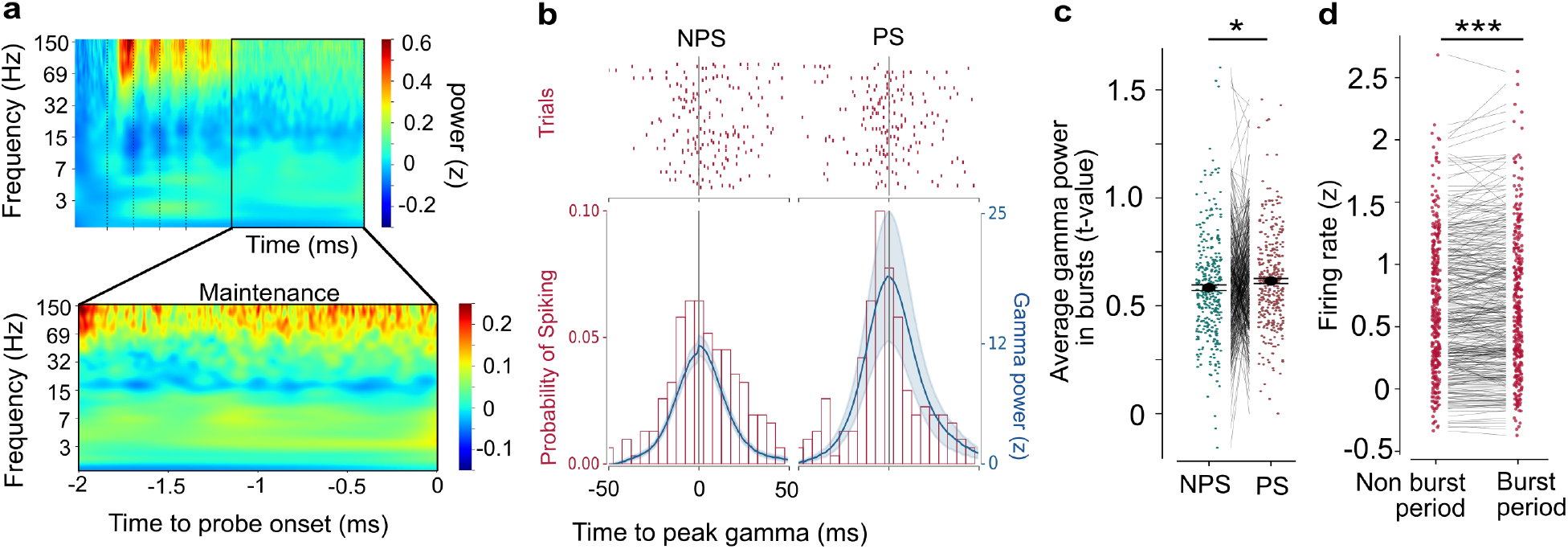
Relationship between gamma and spiking during sequence maintenance. **a**, Average time frequency spectrum (z-scored power relative to baseline) showing increases in gamma power following stimulus onset during encoding period (above) and memory maintenance period (bottom) (N=291). The black dotted lines represent the stimulus onset times during encoding. **b**, A representative neuron exhibiting elevated firing during periods of peak gamma power of each trial during maintenance. The raster plot represents trials in which the NPS (left) and PS (right) were shown. The PSTH (5ms bins) is shown (bottom) with gamma power in blue (mean *±* s.e.m). **c**, Comparison of the magnitude of increase in gamma power between stimulus related bursts identified in channels with responsive neurons. (p *<* 0.05, permutation based t-test of median t-value per channel, N = 340) **d**, Comparison of firing during burst and non-bursts periods (p *<* 0.001, one-sided permutation test, N= 340).

### 2.1 Theta, gamma and spikes are coupled during memory maintenance

Next, we assessed the relationship between theta, gamma and spiking. Similar to previous studies [14, 1], theta power significantly increased during memory maintenance for channels exhibiting stimulus related responses in *gamma* vs. not (U = 75,810, p *<* 0.001, Mann–Whitney U test). In addition, responsive channels showed significantly stronger PAC between theta and gamma (30 to 100 Hz, Tort’s modulation index [24], p *<* 0.001, Mann–Whitney U test; N=108; Fig.3b,c). Theta–gamma coupling was also significantly greater during PS maintenance than NPS (p *<* 0.01, one-sided WSR test ; Fig.3d) suggesting that coupling magnitude depends on encoded stimulus preference. The magnitude of coupling, however, did not vary significantly with the position of the preferred stimulus in the sequence (p *<* 0.05, two-sided Kruskal-Wallis test). However, gamma power was systematically modulated by theta-*phase* as a function of stimulus position (Fig. 3f-g, WSR test, FDR correction; N =141), and this effect was also significantly enhanced for correct vs. incorrect trials (p *<* 0.001, WSR test; Fig. 3h). Accordingly, gamma-theta PAC magnitude was significantly larger for incorrect vs. correct trials when tested *across* all stimulus positions (p *<* 0.05, one-sided WSR test, N = 108). These results support previous studies proposing theta-gamma coupling as a mechanism for organizing multi-item working memory by temporally segregating item-specific representations [11]. Given that both gamma power and SUA encode stimulus position in a phase dependent manner, and exhibit amplitude coupling, we finally asked whether they align to similar phases of the ongoing theta rhythm. Using a subset of LFP-SUA pairs responding to the identical preferred stimulus, we observed no significant difference in their preferred phase angle (N = 41, n.s., one-sided WSR test; Fig. 3i). This was confirmed when analysing the circular correlation coefficient between the preferred theta phases of gamma power and SUA firing, which was significantly higher than phase-shuffled surrogate distributions (r=0.1817, p *<* 0.05, permutation test; N = 41 pairs; Fig. S2d). As expected, preferred gamma phase also revealed no consistent relationship between theta phase order and stimulus sequence order (Fig. S2b-c). In summary, our results confirm earlier findings on increased theta-gamma phase coupling during memory maintenance [1]. Additionally, we firstly demonstrate that gamma activity exhibited robust phase-alignment to theta oscillations in a similar manner as spiking when preferentially responding to the identical stimulus. Our results suggest that gamma and spiking are functionally coupled by theta phase in human MTL, enabling phase-coding of distinct stimuli within a sequence.

**Fig. 3.**
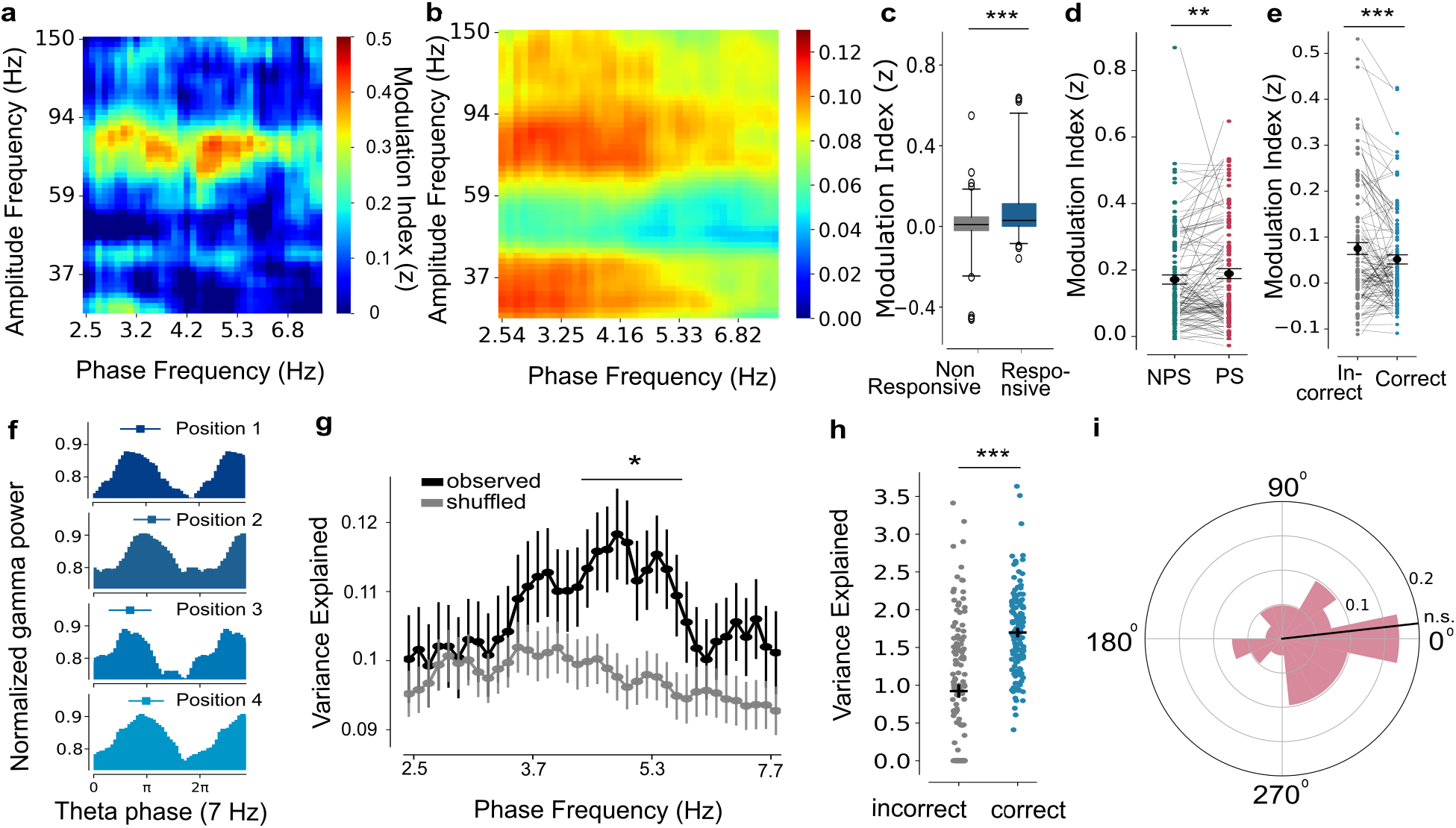
Phase coding and Theta - SUA - Gamma relationship during sequence maintenance. **a**, An example hippocampal channel showing increased Phase amplitude coupling (PAC) in the 60 to 100 Hz range. **b**, Average normalized modulation indices (MI) of all responsive channels (N = 291) **c**, Comparison of PAC magnitude (normalized MI) between non-responsive and responsive channels. p *<* 0.001, Mann Whitney U test. **d**, Pairwise comparison of MI between PS and NPS trials in channels exhibiting significant PAC. p *<* 0.05, one-sided Wilcoxon signed-rank (WSR) test, N = 110 channels **e** Comparison of PAC between correct and incorrect trials in PS trials (p *<* 0.05, one-sided WSR test, N = 108 channels) **f**, Theta phase histograms of gamma power increases (30 - 100 Hz) for the different positions of PS (color coded) for an example channel. Mean preferred phase and circular standard deviation are shown above the histograms in each plot. **g**, Circular variance explained (Vex) between positions for channels exhibiting significant phase coding (p *<* 0.05, Permutation test, N = 141). The average Vex for true (black) and shuffled (gray) position labels. Mean and s.e.m. are plotted (N=141). p *<* 0.05, one-sided WSR test. **h**, Comparison of Vex in correct and incorrect trials for theta frequencies exhibiting highest Vex. (N = 141, p *<* 0.001, one-sided WSR test) **i**, Histogram of the circular mean of position-specific preferred theta phase differences between Gamma and SUA for same PS encoding pairs. The black line represents the circular median (N = 41, n.s. non. significant. p *<* 0.05, one-sided WSR test). The circular correlation co-efficient between the phases is 0.1817 (p *<* 0.05, Permutation test)

## 3 Discussion

We investigated how interactions between spikes, gamma, and theta oscillations coordinate memorized sequences in the human medial temporal lobe (MTL), using simultaneous single-unit and LFP recordings. During encoding, neural firing and gamma activity were temporally coupled. Gamma power encoded stimulus information similar to spiking activity, albeit exhibiting broader stimulus preferences. Interestingly, gamma activity - and stimulus related information it carried - preceded neuronal firing. During memory maintenance, gamma activity was not uniformly enhanced, but organized into short, recurring bursts, that coincided with stronger stimulus-dependent firing. Moreover, both gamma oscillations and single-unit activity were temporally aligned to similar theta phases, but only when encoding similar stimulus content. These findings support a hierarchically nested oscillatory framework spanning slow and fast oscillations that structure spiking to maintain sequential information in human memory.

Encoding-related, stimulus-specific gamma in the human MTL is consistent with previous findings across various memory paradigms and brain regions [9, 10, 17]. The observation that gamma precedes single-unit firing mirrors reports in macaques and suggests that gamma may reflect input from upstream cortical areas, rather than merely indexing local multi-unit activity [20, 3, 4]. Enhanced gamma–spike phase coupling during successful encoding of the preferred stimulus supports this interpretation and aligns with theoretical proposals that precise spike timing—organized into narrow temporal windows—facilitates interregional communication and enhances the representation of behaviourally relevant information [12, 7, 8]. Whether such enhanced coupling directly contributes to information transfer between regions remains an open question [21].

Although overall gamma power did not increase during memory maintenance, it re-emerged in transient bursts throughout the delay period. These bursts were stronger when neurons’ preferred stimuli were part of the maintained sequence and were associated with increased firing. This is in line with prior work suggesting that gamma oscillations dynamically recruit functionally connected cell assemblies to reactivate stimulus-specific ensembles that represent working memory content [6, 17, 22].

Similar to single-unit activity, gamma power was modulated by theta phase. Notably, both gamma activity and spiking were aligned to the same theta phase—but only when encoding similar stimulus information. Consequently, the preferred theta phase of gamma power varied with sequence position, indicating position-specific phase coding. The strength of this phase-specificity was correlated with behavioural performance, underscoring its functional relevance for memory maintenance. However, as with spiking activity, the sequence of gamma-related theta phases did not reflect item order, diverging from a central prediction of a previous conceptual model, which posits that items are represented at successively lagged theta phases [15]. Overall, our results suggest that theta provides a global temporal reference, while gamma subdivides this timescale for item-specific reactivation of cell assembly firing. Future work should explore whether nested theta-gamma-spiking relationships change with cognitive parameters affecting memory - including memory load or item and sequence familiarity. More importantly, it remains to be determined, whether this oscillatory framework also supports local and long-range coordination between different brain regions [23, 5]. In summary, our work adds to growing evidence that interactions between spiking activity and oscillatory dynamics may reflect a core computational principle supporting diverse memory processes in the human brain.

## 4 Methods

For details of the experimental paradigm, recording setup and preprocessing, see [14]; The preprocessing pipeline includes subtraction of average spike waveforms from the raw signal to minimize contamination of local field potentials by action potentials.

### LFP and spike analyses

Time-frequency analysis of LFP signals was performed using complex Morlet wavelets (7 cycles) on data downsampled to 1 kHz. The analytic signal y(t,f) was obtained by convolving the raw LFP time series x(t) with complex wavelets w(t,f) across 150 logarithmically-spaced frequencies (f, 1.5-150 Hz), where f represents the center frequency of each wavelet. Task associated oscillatory power was quantified as z-scores relative to baseline (500 ms pre-stimulus window), calculated by subtracting mean baseline power and dividing by baseline standard deviation for each trial before averaging (Fig. 1b).

To assess stimulus responsiveness in gamma power, we compared pre-stimulus baseline to post-stimulus activity across channels. Baseline was defined as average gamma power within a 500 ms window preceding the first stimulus. Post-stimulus activity was analysed using 100 ms bins with 50 ms overlap from 100-600 ms after stimulus onset. Statistical comparison was performed using one-tailed Wilcoxon signed-rank tests on each time bin versus baseline across all trials of a particular stimulus, independent of position (n= 224 /112 trials). P-values were corrected for multiple comparisons using the Simes procedure (n=13 time windows). We analyzed 1411 units across MTL regions after removing artefactual units identified through semi-manual spike sorting using Waveclus 2.0 and Combinato. By applying the aforementioned statistical procedure to spiking activity, we identified 422 stimulus-responsive units distributed across hippocampus. This approach allowed direct comparison of stimulus response properties between gamma oscillations and neuronal firing within the same MTL regions. A channel or neuron was classified as responsive if there was a significant increase in gamma power or firing rate relative to baseline (p < 0.05, one-sided WSR test) within the region-specific time ranges: 250-600 ms post-stimulus for hippocampal, entorhinal cortex, and amygdala neurons, or 200-550 ms for parahippocampal neurons, accounting for observed regional differences in response latency [18]. This approach allowed direct comparison of stimulus response properties between gamma oscillations and neuronal firing within the same MTL regions.

### Comparison of stimulus encoding properties

We tested how well the gamma power and neuronal firing differentiated between stimuli using two analyses. We first defined selectivity as hedge’s g or the normalized difference between the activity for the most preferred stimulus and second preferred stimulus, defined based on the statistical test p values described above. The formula for hedge’s g is as follows :

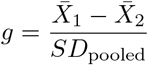

where 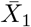 and 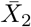are the means of the two groups, and *SD*_pooled_ is the pooled standard deviation.

Stimulus identity decoding was performed separately for gamma power (60–100 Hz) and neuronal firing rates using responsive channels / units identified per session. A multinomial logistic regression classifier was trained to decode stimuli using 101 iterations of stratified shuffle-split cross-validation (test size: 15%, training size: 85%), maintaining class balance across splits and assessing model performance via accuracy (proportion of correct predictions). Shuffled distributions were generated by permuting stimulus labels and decoding performance was compared.

### Precedence of gamma power increase

Response latencies of SUA and gamma were estimated to compare the temporal precedence of gamma power and neuronal firing rate increases following their respective preferred stimulus onset. Gamma power (60–100 Hz) and firing rates were computed on single trials, and rise time was defined as the latency to reach 50% of the peak response after the first 100 ms following stimulus onset, accounting for expected delays in information transmission to the MTL regions [18]. Statistical comparisons were performed on median rise time across trials in a channel or neuron. To verify the robustness of response latencies, we repeated the analysis using different smoothing parameters (15 to 40 ms) for firing rate estimation. In all cases, gamma power increase consistently preceded SUA increase (p *<* 0.05, WRS test) To assess the temporal relationship between gamma power and neuronal firing rates, we computed cross-correlations using trial-averaged signals for each neuron and channel. Gamma power (60–100 Hz) and firing rates were extracted per trial, aligned to PS of neuron onset, and averaged across trials. Time segments for analysis were defined as 200 ms windows centered on the peak of neuronal firing rates. For each pair, Spearman correlation coefficients were calculated across lags from -75 ms to +75 ms, with negative lags indicating gamma leading firing. For each neuron, the peak correlation and corresponding lag were identified.

### Spike Phase coupling analysis

The following procedure was used to quantify Spike Phase Coupling (SPC) and assess differences in SPC between experimental epochs. Phases computed using complex Morlet’s wavelets explained above were extracted during times at which spikes occurred recorded, in the same microwire for frequencies in the gamma range (60 (30) - 100 Hz for encoding/maintenance, respectively) based on the distribution of peak gamma frequencies during the two task periods (Fig. S1b). Phases that occurred at the time window corresponding to encoding period (same window used for defining responsive units, see LFP and spike analyses) were retained. Spike-LFP pairs were subjected to a Rayleigh test to check for non-uniformity of phase distributions during the entire encoding or delay without an bias towards PS or NPS. Neurons that passed the Rayleigh’s test for uniformity and had at least 10 spikes were included for comparison of spike phase coupling between PS and NPS trials. In this step, Mean Vector length was calculated during encoding for trials with PS and NPS separately. To avoid biases in mean vector length due to spike counts, we equalized the number of spikes in each condition by randomly sampling 10 spikes across conditions in each spike-lfp pair and computing the mean vector length. This step was repeated 500 times and the average mean vector length was taken as the spike phase coupling magnitude to a particular frequency. The mean vector length for the gamma frequency range (60 - 100 Hz) were averaged to obtain the spike phase coupling to the gamma band.

### Phase Amplitude Coupling (PAC) analysis

To determine if gamma amplitude increase is dependent on theta phases, we used modulation index (MI) as our phase amplitude coupling metric [24]. The method quantifies how much a distribution of gamma amplitudes over phase bins deviates from a uniform distribution. We calculated PAC for each combination of amplitude between gamma frequencies between 30 - 100 hz and phases between theta frequencies 2.5 - 8 Hz during the entire delay period (2000 ms) in 500 ms windows, z-scored the MI values using baseline PAC and averaged across to obtain the PAC comodulogram per channel. Among the responsive channels, we identified the channels with significant phase amplitude coupling using cluster-based permutation tests. Channels with at least one significant cluster spanning across were considered to be significant and called PAC+ channels. We identified 108 PAC+ across the MTL regions. Among these channels, we assessed the difference between PS and NPS by computing the PAC comodulograms separately for trials in which PS and NPS were shown. To assess relationship to behaviour, we computed the comodulograms separately for the correct and incorrect trials among trials during which PS was shown.

### Identification of gamma bursts

To identify periods of increased gamma power or bursts associated with a particular stimulus during the memory maintenance period, we employed a cluster-based permutation testing approach. Time-resolved gamma power was first computed for trials with the stimulus and compared to trials without using t-values as the permutation statistic. To minimize contamination from effects not related to the task, trials with peaks in power exceeding four times the z-scored values during maintenance period in the raw data were excluded from the gamma burst analysis. Temporally contiguous time points exceeding a predefined significance threshold (p *<*0.05, independent samples t-test) were grouped into clusters. For each cluster, the t-values were summed to yield a cluster-level statistic. To assess the significance of these clusters, we generated a null distribution by randomly shuffling the conditions 199 times and computing the maximum sum of clusters for each permutation. Observed clusters were considered significant if their test statistic exceeded the 95th percentile of the null distribution. Only the cluster with the maximum t-value exceeding this threshold was retained for further analyses.

### Comparison of Gamma bursts and SUA during memory maintenance

To examine the relationship between stimulus-specific gamma bursts and single-unit activity (SUA) during the memory maintenance period, we first identified time points showing gamma bursts (30–100 Hz) associated with PS or NPS of a neuron. We then extracted spike counts within and outside these burst-associated windows for PS trials. For comparison, spike counts were also extracted from non-preferred stimulus (NPS) trials using NPS specific bursts. To control for potential confounds due to non stimulus specific bursts, we performed a shuffling procedure in which burst-associated time points were circularly shifted within the delay period, preserving the temporal structure and duration of burst and non-burst intervals. For comparison, average spike counts were also extracted from PS trials using multiple shuffled time windows (N = 999 shuffled). This approach allowed us to test whether gamma bursts associated with PS elicited stimulus-specific increases in SUA and compare with overall maintenance firing rate. We repeated the analyses by identifying the gamma bursts associated with the non preferred stimulus specific bursts of the neuron to confirm if the neuron selectively modulate its firing only for the preferred stimulus. We only included neurons that were significantly responsive to one or two stimuli to test stimulus specific spike association with gamma burst windows.

For spike–phase coupling analyses, we computed the phase of the gamma oscillation at spike times separately for spikes occurring within and outside of burst-associated windows and computed average mean vector length accounting for spike count differences as described above (spike phase coupling analyses). This enabled assessment of whether SUA exhibited stronger phase-locking to gamma oscillations specifically during stimulus-relevant burst events.

### Phase coding of stimulus position analyses

To assess whether the preferred theta phase of gamma power during the memory delay was modulated by stimulus position, we computed trial-wise estimates of the mean phase of gamma power for the preferred stimulus of a channel. For each trial, average gamma power (30 - 100 Hz) across the entire delay period was binned according to the theta phase bins (36 bins) and a von mises distribution was fit to get the Preferred phase of gamma power per trial. This was repeated for each theta frequency between 2.5 to 8 Hz. We subsequently computed the circular variance explained between different experimental conditions (in our case stimulus positions 1–4) by quantifying the ratio of variance within conditions relative to the variance across conditions ([14]). To assess phase ordering across positions, we centred phase estimates by subtracting the circular mean phase across all positions from each position’s phase. Phases were sorted and the stimulus position order corresponding to the sorted phase order were obtained.

### Gamma and SUA phase alignment

To examine the alignment of gamma power and SUA firing to theta phases in a stimulus-specific manner, we analysed “same preferred stimulus” (PS) encoding pairs from the same channel that exhibited significant phase coding in gamma power. For each pair, we identified the theta frequency at which gamma power showed maximal phase separation and extracted the preferred theta phase for both gamma power and SUA firing.

First, we calculated the circular mean of the position-specific preferred theta phase differences between gamma power and SUA for each pair, and tested whether this distribution differed significantly from zero using a one-sided Wilcoxon signed-rank test. Next, we computed the circular correlation coefficient between the preferred theta phases of gamma power and SUA firing. Statistical significance was determined by comparing the observed correlation to a distribution generated from 9,999 phase-shuffled surrogates. Only units from channels with significant phase coding were included in the above analyses.

## 5 Supplementary information

**S1.**
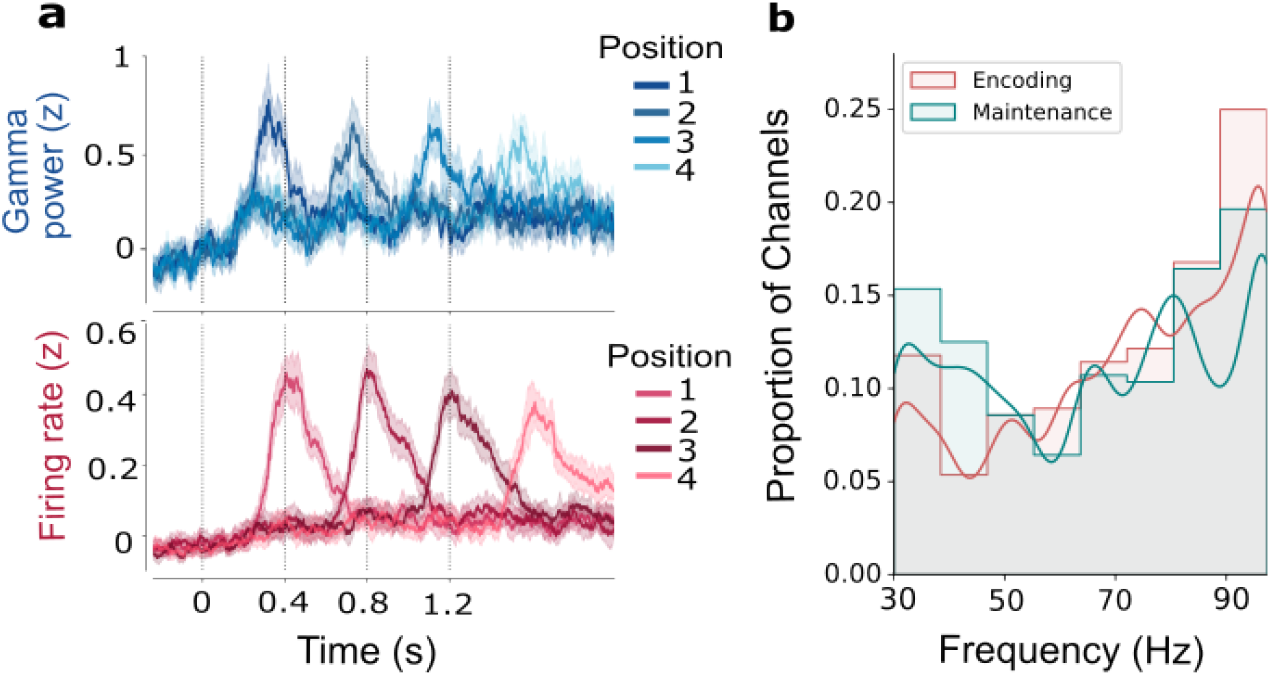
Gamma power increases following stimulus onset during encoding. **a**, Median and 95% C.I. are shown for z-scored gamma power (60 - 100 Hz) and firing rates following PS onset at different positions (colour coded). **b**, Distribution of peak frequencies in responsive channels during encoding and memory maintenance (N = 291), identified based on median power in each period. The maintenance period exhibits a relative increase in the proportion of channels with peak frequencies in the 30–60 Hz range. Analyses during maintenance were performed across the entire 30–100 Hz range.The distribution of peak frequencies deviated significantly from uniformity during encoding (Kolmogorov–Smirnov test, p *<* 0.001) and to a lesser extent during the maintenance delay (p *<* 0.05).

**S2.**
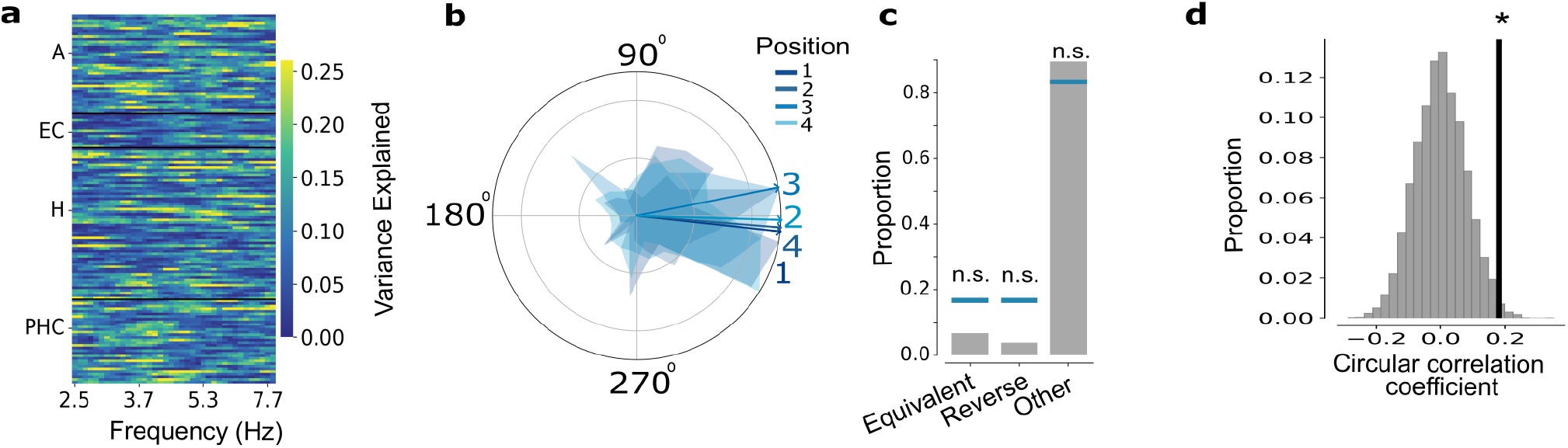
Preferred phase of gamma power increase varies with stimulus position. **a**,Circular variance explained (Vex) between positions for channels exhibiting significant phase coding (N = 141, p *<* 0.05, Permutation test). **b**,Histogram of preferred theta phase and mean direction of gamma power increases across channels shown in **b** per stimulus position (colour coded). Circular differences in phase with respect to mean phase across all positions per unit. The numbers correspond to the position to visualize the order of mean direction better. **c**, The proportion of channels for which the phase order is equivalent to the item order, or reversed or is different. The blue lines denote the proportion expected by chance (1/6, 1/6, 5/6, respectively) n.s. non-significant z-test for proportions. **d**, Observed circular correlation (black line) co-efficient between position specific theta preferred phases of Gamma and SUA for pairs with same PS (N = 41 pairs) compared to the shuffled surrogates (p *<* 0.05, Permutation test)

## Declarations

## Acknowledgements

This work was supported by the German Research Foundation (DFG): Priority Programme SPP 2411 (PN 520287829), SFB 1233 (PN 276693517) and the Excellence Cluster “Machine Learning in Science” Tuebingen (EXC number 2064/1 – PN390727645). MJP is supported by International Max Planck Research School for The Mechanisms of Mental Function and Dysfunction. SL is supported by the Clinician Scientist program of the Medical Faculty Tübingen funded by (DFG, PN 493665037). We thank Matthijs Pals for comments on the manuscript.

## Ethics approval and consent to participate

The study was approved by the Medical Institutional Review Board of the University of Bonn (accession number 095/10 for single-unit recordings in humans in general and 249/11 for the current paradigm in particular) and adhered to the guidelines of the Declaration of Helsinki.

## Author contribution

M.J.P. designed, implemented and performed formal analyses and visualization. J.N. and F.M. designed the experiment, collected data. J.N. implemented the experiment. R.S. provided administrative support. S.L. and J.N. curated data. S.L. conceived the study, designed analyses and supervised and administered the project. M.J.P. and S.L. wrote the paper. All authors reviewed the results and approved the final version of the manuscript.

